# The Arabidopsis autophagy cargo receptors ATI1 and ATI2 bind to ATG8 via intrinsically disordered regions and are post-translationally modified upon ATG8 interaction

**DOI:** 10.1101/406538

**Authors:** Ida Marie Zobbe Sjøgaard, Simon Bressendorff, Andreas Prestel, Swathi Kausika, Emilie Oksbjerg, Birthe B. Kragelund, Peter Brodersen

## Abstract

Selective autophagy has emerged as an important mechanism by which eukaryotic cells control the abundance of specific proteins. This mechanism relies on cargo recruitment to autophagosomes by receptors that bind to both the ubiquitin-like AUTOPHAGY8 (ATG8) protein through ATG8 interacting motifs (AIMs) and to the cargo to be degraded. In plants, two autophagy cargo receptors, ATG8 Interacting Protein 1 (ATI1) and 2 (ATI2), were identified early on, but their molecular properties remain poorly understood. Here, we show that ATI1 and ATI2 are transmembrane proteins with long N-terminal intrinsically disordered regions (IDRs). The N-terminal IDRs contain the functional AIMs, and we use nuclear magnetic resonance spectroscopy to directly observe the disorder-order transition of the AIM upon ATG8 binding. Our analyses also show that the IDRs of ATI1 and ATI2 are not equivalent, because ATI2 has properties of a fully disordered polypeptide, while ATI1 has properties consistent with a collapsed pre-molten globule-like conformation Interestingly, wild type ATI1 and ATI2 exist as distinct post-translationally modified forms. Specifically, different forms are detectable upon mutation of the AIM, suggesting that interaction of ATI1 and ATI2 to ATG8 is coupled to a change in their post-translational modification.

## INTRODUCTION

Autophagy is a major pathway of macromolecular degradation in eukaryotes. It involves sequestering of cytoplasmic material by a growing double-membrane structure, the isolation membrane or phagophore. Membrane assembly at the phagophore is critically dependent on a ubiquitin-like protein, AUTOPHAGY8 (ATG8), that is activated by conjugation to phosphatidylethanolamine at its carboxy terminus [1]. Upon maturation to double-membrane vesicles, the so-called autophagosomes traffic to fuse with membranes surrounding lytic compartments, in plants the central vacuole, for degradation of the inner vesicle and its contents [1]. While autophagy has long been recognized as a major pathway of degradation and recycling of bulk cytoplasmic material, progress during the last 10 years has revealed that it can also be employed for degradation of specific targets, including individual proteins and protein complexes [2,3]. Such selective autophagy involves cargo receptor proteins that contain one or more ATG8-interacting motifs (AIMs), short linear motifs (SLiMs) with the consensus sequence Trp/Phe/Tyr-X-X-Leu/Ile/Val (W/F/Y-X-X-L/I/V) [4-6], and additional interaction sites for specific cellular proteins. This combination of functional elements enables cargo receptors to act as adaptors to recruit specific proteins to the phagophore. Not all proteins containing the W/F/Y-X-X-L/I/V consensus SLiM bind to ATG8, however. It is likely that the structural context of the AIM is critical to its function, because a recent examination of many *bona fide* AIMs proven experimentally to confer ATG8 binding showed that similar to most SLiMs, they tend to occur in parts of proteins predicted to be structurally disordered [7]. Nonetheless, few studies have addressed the structural context of functional AIMs by direct experimentation.

The mammalian p62 and NBR1 proteins were among the first AIM-containing cargo receptors to be identified. They interact with ubiquitylated target proteins or protein aggregates and mediate the degradation of such targets via autophagy [8-10]. A mixed p62/NBR1 homologue has also been identified in Arabidopsis (AtNBR1) [11], and has recently been shown to play a role in selective autophagy of the cauliflower mosaic virus (CaMV) capsid protein P4 and viral particles, thereby limiting infection by CaMV [12]. Similarly, entire plant proteasomes can be degraded by selective autophagy dependent on the AIM-containing subunit RPN10 [13]. Indeed, the recent identification of a series of other plant cargoes and cargo receptors has made it clear that selective autophagy is widespread and plays fundamental roles in aspects of plant biology related to stress adaptation. For example, the abundance of the transcription factor BES1 essential for brassinosteroid-dependent growth promotion is controlled by selective autophagy in response to stress via the cargo receptor DSK2 [14]. Selective autophagy can also be employed to shape the duration of protein induction in response to stress. For example, transient rather than lasting induction of the stress-related membrane protein TSPO by abscisic acid is ensured by selective autophagy dependent on intact AIMs in TSPO itself [15]. Such regulated exposure of AIMs in response to stimuli can be controlled by post-translational modification of cargoes as illustrated by a recent study of the Arabidopsis enzyme *S-*nitrosoglutathione reductase (GSNOR), important for the biology of reactive nitrogen species. *S-*nitrosylation of GSNOR upon hypoxia induces conformational changes that expose AIMs in GSNOR, thereby leading to its degradation via selective autophagy [16].

In addition to these examples of autophagy cargoes and cargo receptors, the Arabidopsis <u>AT</u>G8-<u>I</u>nteracting proteins ATI1 and ATI2 represent two likely, but atypical autophagy cargo receptors: Like other cargo receptors, they contain candidate AIMs, and they interact with the ATG8 isoform ATG8f in yeast two-hybrid assays and *in planta* [17]. They also colocalize with ATG8 [17], and ATI1 may deliver select plastidial cargo to vacuolar degradation, because it colocalizes with plastid-associated bodies and is required for autophagy-dependent turnover of a stromal protein [18]. Interestingly, however, ATI1 and ATI2 are only rarely found with ATG8 in autophagosomes, but localize mainly to ER-derived vesicles that traffic to the vacuole [17]. ATI1 and ATI2 function may therefore involve ATG8-dependent vacuolar proteolysis via a vacuolar trafficking pathway that does not involve autophagosomes. Despite these interesting features of ATI1 and ATI2, precise molecular identities of cargoes have not been elucidated, and basic molecular properties of the ATI1 and ATI2 proteins remain unclear: it is not known whether the proteins are transmembrane or peripheral membrane proteins, the precise site of ATG8 interaction is unknown, and there is confusion as to whether the N-terminal part of ATI1 is globular or intrinsically disordered. This has serious ramifications for a precise understanding of their molecular function, because globular and intrinsically disordered proteins have fundamentally different properties not only structurally, but also functionally, for example as mediators of allosteric effects [19] that could control stimulus-dependent degradation of cargo proteins. In addition, information on their potential post-translational modification is missing.

Here, we use a series of biochemical and biophysical experiments to show that ATI1 and ATI2 are transmembrane proteins, and that they contain long disordered N-termini with the functional AIM close to the N-terminus. In both cases, nuclear magnetic resonance (NMR) spectra provide direct evidence that tryptophan residues contained in the AIMs are placed in highly disordered regions. The N-terminal IDRs differ between the two proteins, however, because ATI1 has molten globule-like properties, while ATI2 has properties of a fully disordered polypeptide. Interestingly, ATI1 and ATI2 are post-translationally modified *in vivo* depending on an intact AIM, suggesting that ATG8 interaction is linked to their post-translational modification.

## RESULTS

### ATI1 and ATI2 are transmembrane proteins

Although ATI1 and ATI2 are associated with the endoplasmic reticulum (ER) or with ER vesicles *in vivo* [17], it remains unclear whether they are transmembrane or peripheral membrane proteins. The Hidden Markov Model-based predictor TM-HMM [20] predicts a single transmembrane helix in ATI1 such that the 180-residue N-terminal domain is cytoplasmic while the 56-residue C-terminal part is luminal (Fig. 1A). On the other hand, ATI2 is not predicted to contain a transmembrane helix despite the fact that the sequence of the putative transmembrane helix in ATI1 is closely related to an analogously located sequence segment in ATI2 (Fig. 1A). To test experimentally whether ATI1 and ATI2 are transmembrane or peripheral membrane proteins, we constructed stable transgenic Arabidopsis lines expressing N-terminally 3xHA-tagged versions. Both ATI1 and ATI2 were found exclusively in the insoluble fraction (Fig. 1B). Microsomal fractions were then washed with either high salt (1M KCl) or alkaline solution (0.1 M Na_2_CO_3_, pH 10) to extract peripheral membrane proteins, or with 1% deoxycholate to solubilize membranes, including transmembrane proteins. ATI1 and ATI2 remained in the insoluble fraction upon high salt and pH washes, but were readily dissolved in 1% deoxycholate. Thus, ATI1 and ATI2 are transmembrane proteins.

**Figure 1:**
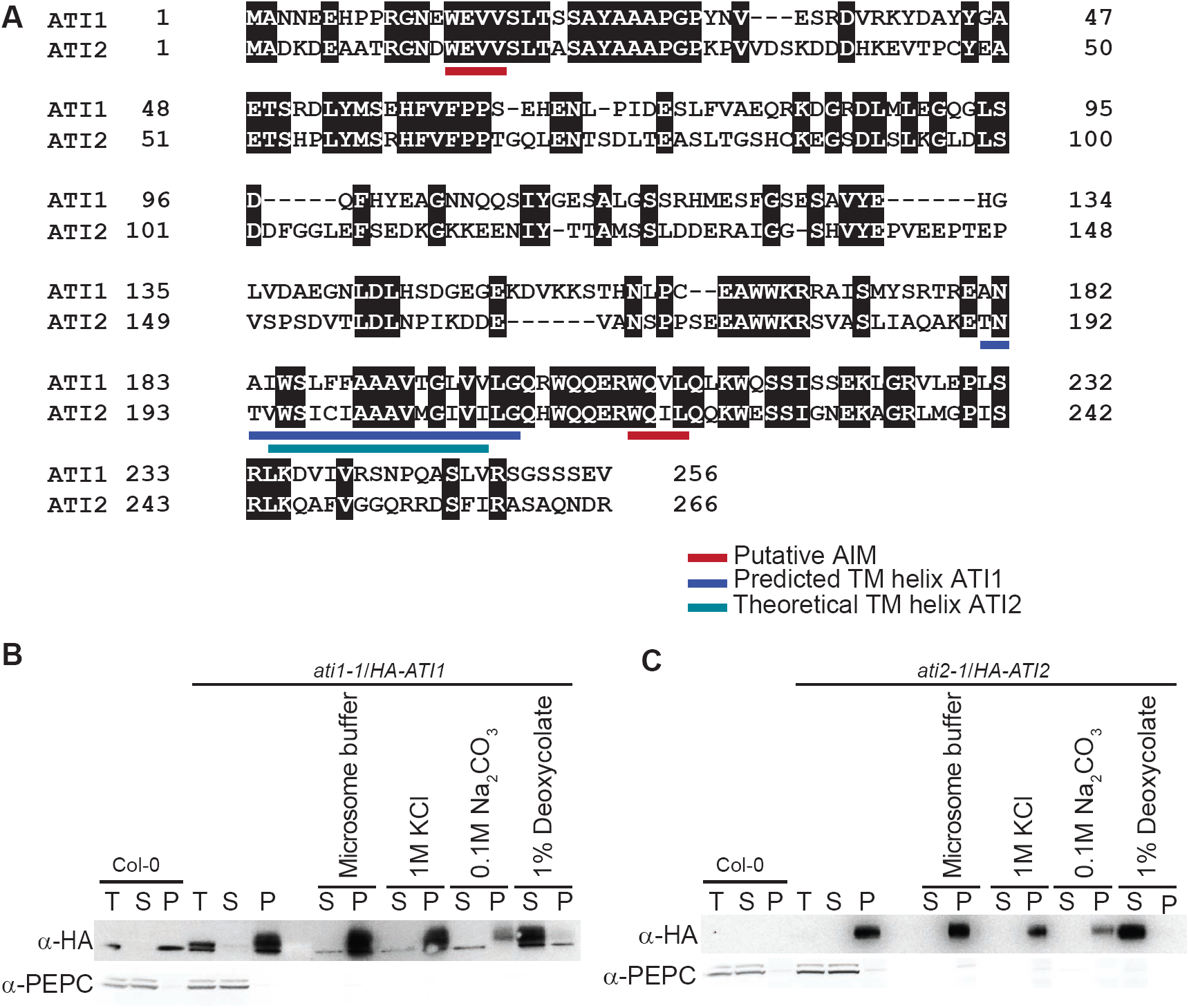
ATI1 and ATI2 contain a transmembrane domain. **A.** Boxshade representation of an alignment of ATI1 and ATI2 (Emboss Needle Pairwise Sequence Alignment) [47] highlighting the TM-HMM-predicted transmembrane helix of ATI1 (181-200) with a blue bar. The analogous region in ATI2, not predicted by TM-HMM to be a transmembrane helix, is highlighted with a cyan bar. The putative N-terminal and C-terminal AIMs are highlighted with red bars. **B.** Microsomal fractions isolated from Col-0 and *ati1-1/HA-ATI1* seedling tissue treated with either micro-some buffer, 1 M KCl, 0.1 M Na_2_CO_3_, or 1% sodium deoxycholate for 1 hr at 4°C. Membranes were then pelleted by 100.000 rcf centrifugation and soluble (S) and pellet (P) fractions were subjected to SDS-PAGE and immunoblotting with HA antibodies and with antibodies recognizing the abundant soluble enzyme phosphoenolpyruvate carboxylase (PEPC). **C.** Same analysis as described in B, but performed with samples prepared from seedlings of Col-0 and *ati2-1/HA-ATI2*.

### Conflicting predictions on intrinsic disorder in N-terminal regions of ATI1 and ATI2

To get a first understanding of protein properties of ATI1 and ATI2, we used the prediction programs IUPred2A [21-23], Globplot [24] and PONDR [25-27] to predict any regions of intrinsic disorder. For ATI2, all predictions converged on the conclusion that the N-terminal 190 residues are intrinsically disordered while some of the C-terminal 70 residues are structured (Fig. 2B). For ATI1, on the other hand, the IUPred and Globplot predictors favour a globular fold of the N-terminal part while PONDR indicates high likelihood of intrinsic disorder (Fig. 2A). We therefore set out to determine experimentally whether the N-terminal ∼180 residues of ATI1 (M1-E180; ATI1-N) and ATI2 (M1-T193; ATI2-N) are intrinsically disordered or not.

**Figure 2:**
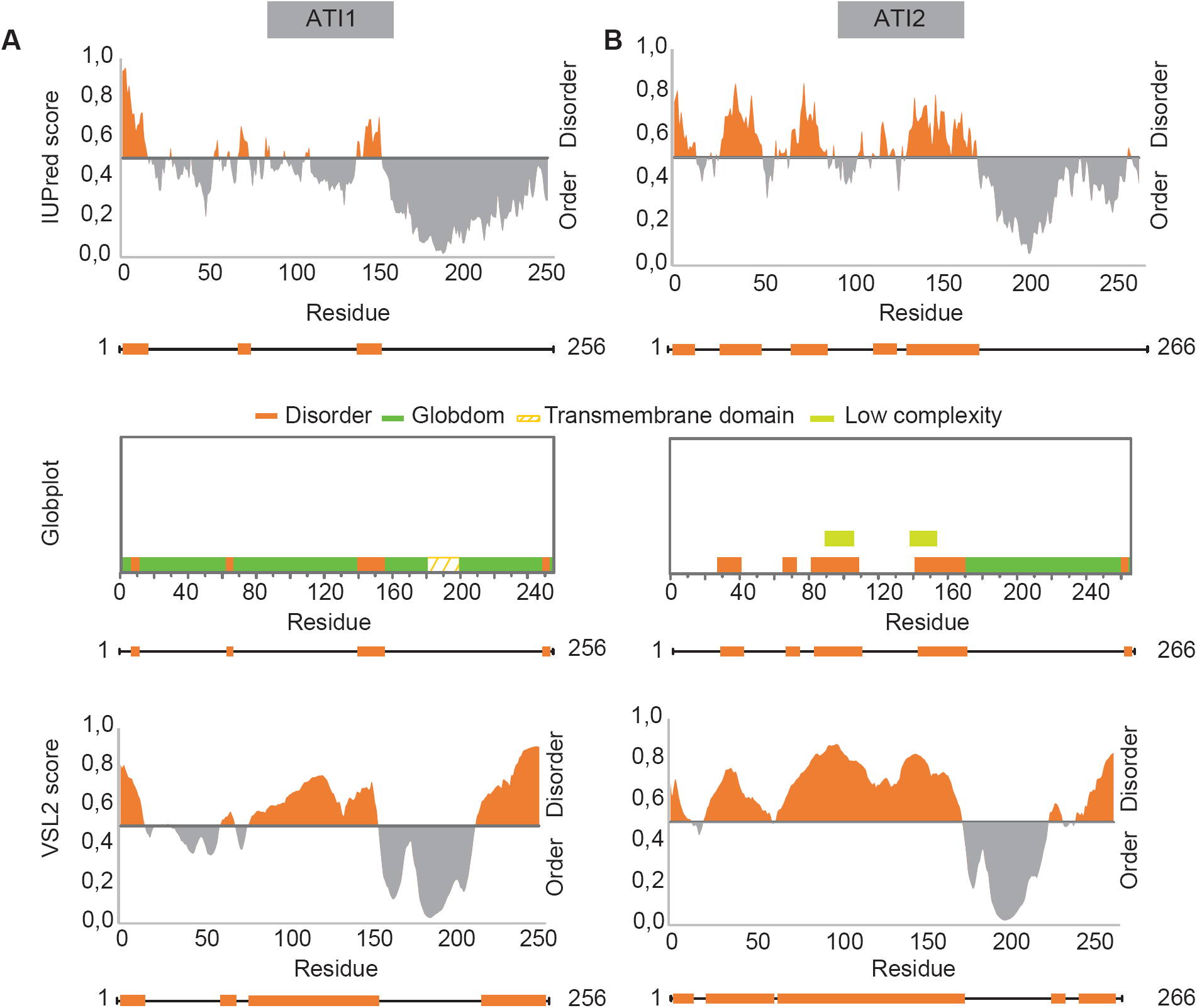
Predictions of intrinsic disorder for ATI1 and ATI2. Regions of intrinsic disorder of ATI1 (**A**) and ATI2 (**B**) predicted by IUPred2A (first panel), Globplot (second panel) and PONDR (VSL2, third panel) using the primary structure as input and shown as a schematic diagram below each output with orange boxes indicating disorder.

### ATI1-N and ATI2-N exhibit large Stoke’s radii, slow migration in SDS gels and no tendency to aggregate at elevated temperatures

We first expressed ATI1-N and ATI2-N in *E. coli* as glutathione-*S*-transferase (GST) fusions with tobacco etch virus (TEV) protease sites engineered to liberate the exact ATI1-N and ATI2-N protein sequences upon cleavage with TEV protease. The proteins were purified by glutathione affinity chromatography, followed by cleavage with His_6_-TEV, and size-exclusion chromatography. We noticed that the migration rate of ATI1-N and ATI2-N in SDS-PAGE was slower than expected by their molecular weights (Mw_ATI1-N_ = 20.46 kDa; Mw_ATI2-N_ = 21.02 kDa) (Fig 3A). This simple observation represents a first indication that ATI1-N and ATI2-N may be intrinsically disordered, because IDRs are typically depleted in hydrophobic residues, and, consequently, tend to bind less SDS, explaining their abnormally slow mobility in SDS-PAGE. We next subjected protein samples of ATI1-N, ATI2-N and the similarly sized, globular Prolactin (PRL, 23 kDa) to size exclusion chromatography (Fig. 3B) in which the elution volume decreases with the Stoke’s radius of the protein, such that folded proteins elute at larger volumes than IDPs of similar molecular weight. ATI2-N elutes at the smallest volume (67 ml), followed by ATI1-N (75 ml), and finally the globular PRL (85 ml), suggesting that the Stoke’s radii (r) increase in the order r(PRL) < r(ATI1-N) < r(ATI2-N). Since we cannot exclude that ATI1-N and ATI2-N form folded dimers, we subjected ATI1-N, ATI2-N and the globular GST to heat treatments for 10 min at 75°C or 5 min at 95°C, and separated aggregated from soluble protein by centrifugation (Fig. 3C-D). These aggregation tests showed that, in contrast to the GST control, ATI1-N and ATI2-N remained soluble upon subjection to elevated temperatures that unfold most globular proteins from non-thermophilic organisms. These observations of behavior in SDS-PAGE, in size exclusion chromatography and upon heat treatment are consistent with ATI1-N and ATI2-N being intrinsically disordered, but do not constitute definitive proof of this on their own.

**Figure 3:**
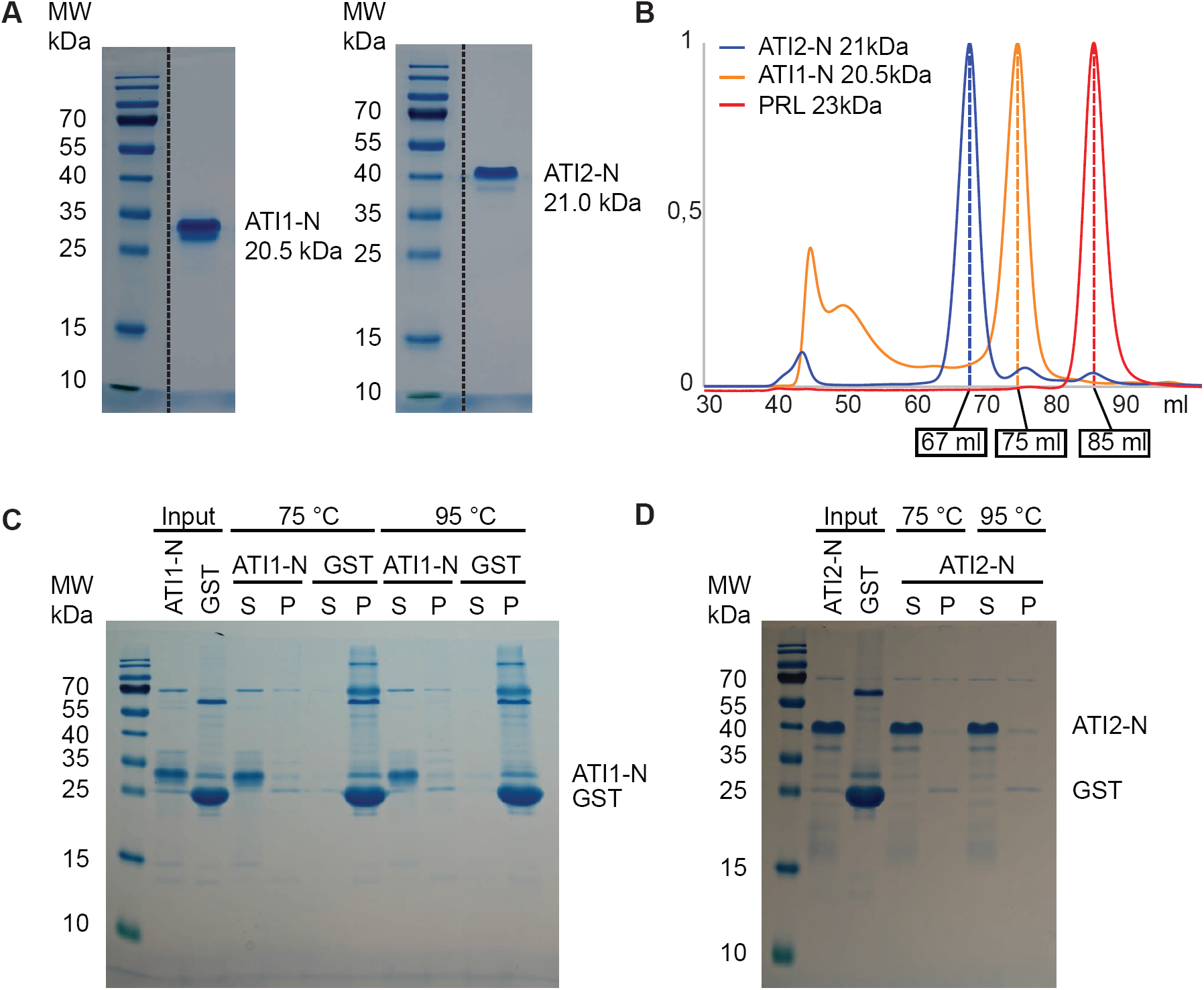
Behaviour of ATI1-N and ATI2-N in SDS-PAGE, in size exclusion chromatography and at elevated temperatures. **A.** Migration of ATI1-N (20.5 kDa) and ATI2-N (21.0 kDa) on a denaturing 14 % polyacrylamide gel. **B.** Elution profile of ATI1-N (orange), ATI2-N (blue) and the globular PRL (red) obtained from size exclusion chromatography. Absorbance is normalized to the peak corresponding to the elution of each of the proteins. Elution volumes are indicated in boxes. **C, D.** Aggregation test of ATI1-N (**C**) and ATI2-N (**D**) when heated for 10 min at 75°C or for 5 min at 95°C. Samples were subsequently centrifuged and supernatants (S) and pellets (P) were analysed by SDS PAGE. GST is included as a folded protein control.

### Circular dichroism spectroscopy indicates random coil properties of ATI1-N and ATI2-N

Circular dichroism (CD) spectroscopy of proteins can provide information on their content and type of secondary structures. In the far-UV region of the spectrum (*λ* < 260 nm), peptide bonds of *α*-helices show diagnostic double minima at 222 nm and 208 nm and a single positive maximum at 190 nm, whereas *β*-sheets generate a single negative minimum around 220-210 nm and a positive maximum at 195 nm [28-30]. In contrast, random coils and disordered conformations are identified by a pronounced negative minimum around 197 nm [31]. To substantiate the bioinformatic and initial experimental analyses described above, we therefore obtained far-UV CD spectra of ATI1-N and ATI2-N. The CD spectra of both ATI1-N and ATI2-N showed characteristic negative minima around 200 nm, indicating a low content of secondary structure (Fig. 4A). In addition, the broad minima at ∼220 nm may indicate some *α*-helical content, in agreement with the secondary structure prediction obtained utilizing the Psipred tool (Fig. 4B) [32,33]. Thus, CD spectroscopy substantiates that both ATI1-N and ATI2-N are largely disordered, and that they may contain *α*-helical segments. We note, however, that although no obvious differences were observed between ATI1-N and ATI2-N by CD spectroscopy, our analyses do not rule out that ATI1-N is more compact as suggested by the results of size exclusion chromatography.

**Figure 4:**
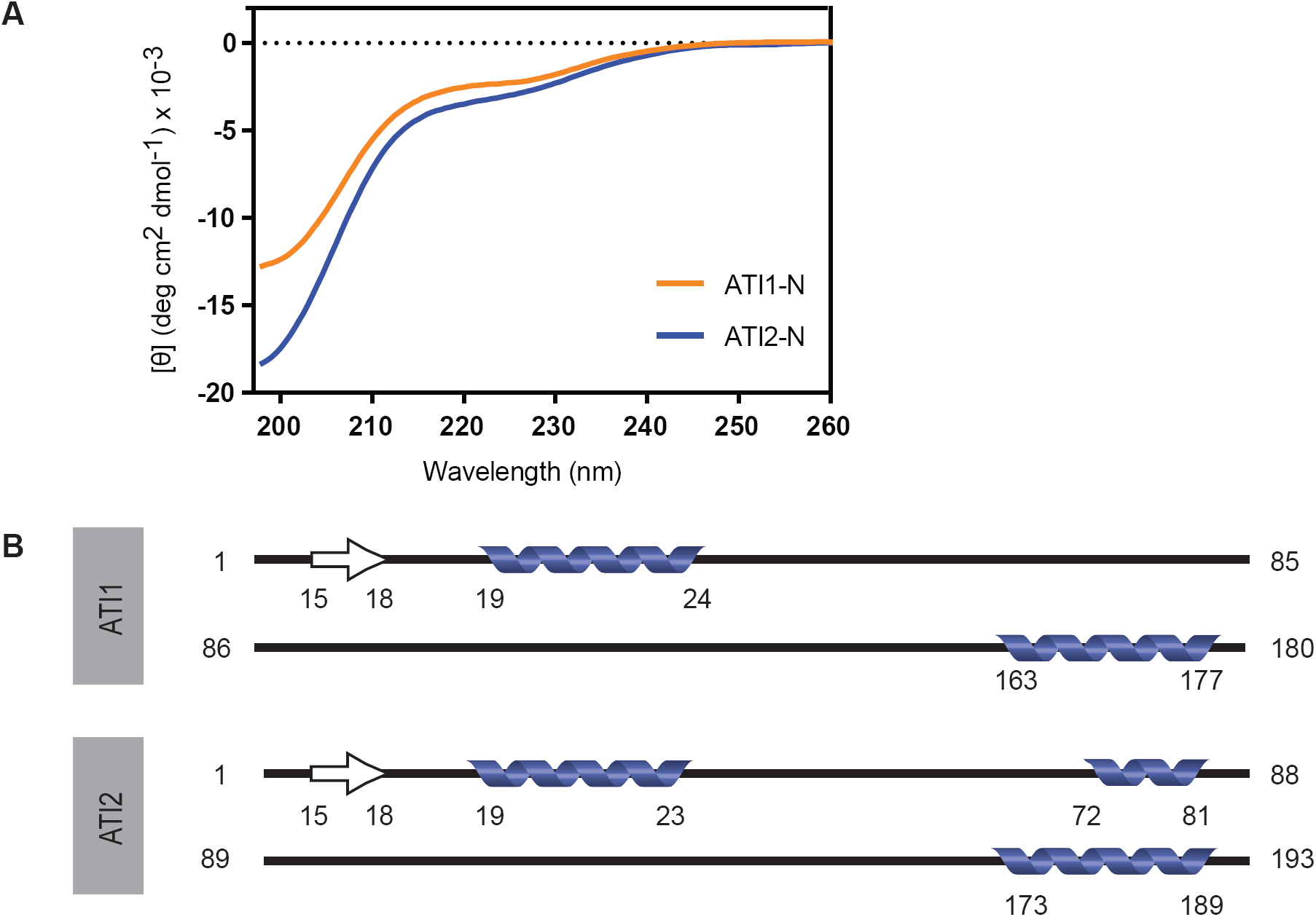
ATI1-N and ATI2-N show a highly disordered conformation in circular dichroism spectroscopy. **A.** Far-UV CD spectra of ATI1-N and ATI2-N characterized by a broad minimum around 222 nm and a deep minimum at 200 nm. Data is given as molar ellipticity with the unit deg cm^2^ dmol^-1^. **B.** Schematics of the secondary structure of ATI1-N and ATI2-N predicted by Psipred. Positions of predicted α-helices (blue helices) and β-sheet (white arrows) are indicated.

### ATI2-N is fully disordered while ATI1-N has molten globule-like properties

To obtain more definitive structural information on ATI1-N and ATI2-N, we performed nuclear magnetic resonance (NMR) spectroscopy of the purified protein samples. Even without assignment of individual peaks to nuclei in specific amino acid residues, one-dimensional (1D) ^1^H NMR spectra can provide information as to whether a protein is folded or disordered. Two effects make this possible. First, the folding of globular proteins generates many different, well-defined chemical environments for both backbone and side chain atoms such that the chemical shifts have a wider distribution in folded proteins than in IDPs. Second, in contrast to the situation in larger folded proteins (>15kDa), nuclei in IDPs give rise to sharp peaks, because of the dynamics leading to longer spin-spin relaxation times. Three regions of 1D ^1^H NMR spectra are particularly informative in these respects:

1. The region of chemical shifts (δ), δ < 1.0 ppm: The methyl groups in alanine, threonine, valine, isoleucine, and leucine sidechains give rise to distinct sharp peaks around 1 ppm in disordered proteins, while the chemical shifts of methyl group show a wider distribution in globular proteins such that signals appear in the otherwise unpopulated part of the spectrum at chemical shifts below 1 ppm [34].
2. The region of chemical shifts close to, but larger than 8.7 ppm: The signals of backbone amide proton signals of IDPs collapse in the interval 8.0 < δ < 8.7 ppm, while their wider distribution in globular proteins gives rise to amide proton signal beyond these boundaries. This effect is detectable at δ > 8.7 ppm where few other nuclei resonate [34].
3. The region of chemical shifts, δ > 10 ppm: The indole NH proton of tryptophan side chains generally give well resolved signals in this region, and can be easily followed in e.g. binding experiments or when changing the environment.

We therefore recorded 1D ^1^H NMR spectra of unlabeled ATI1-N and ATI2-N, and analyzed these three regions of the spectra in detail. ATI2-N shows spectral properties of an IDP in all three regions: methyl protons show a narrow dispersion of chemicals shifts around 0.8 ppm, amide protons resonate at chemical shifts within the interval 8.0 < δ < 8.7 ppm, and the three tryptophan NHs in ATI2-N give rise to sharp peaks reflecting high conformational flexibility in these regions (Fig. 5). Thus, together with the size exclusion chromatography results, we conclude that ATI2-N is a highly disordered IDP with little or no long-range contacts. For ATI1-N, the results are less clear cut. Although most amide and methyl proton signals collapse, characteristic of IDPs and different from fully folded regions, a broader dispersion of signals both in the methyl region and in the amide region was observed compared to ATI2-N (Fig. 5). In addition, the signals of two of the three observable indole NHs of ATI1-N showed a severely increased linewidth (Fig. 5), probably due to a combination of two different contributions: (i) a longer rotational correlation time of residues in structured regions tumbling more slowly than fully unstructured regions with very short local rotational correlation times; (ii) intermediate timescale-exchange between different conformers in an ensemble of conformations. The latter is often observed in disordered regions with transient structure formation and/or transient long-range contacts, consistent with the more compact nature of ATI1-N compared to ATI2-N as suggested by the size exclusion chromatography results. We conclude from the biochemical and biophysical analyses of ATI1-N and ATI2-N that both proteins have characteristics of IDRs, but that important differences between the two are detectable: While ATI2-N behaves like a fully disordered IDR, ATI1-N has properties of a collapsed pre-molten globule-like structure with very low content of secondary structure.

**Figure 5:**
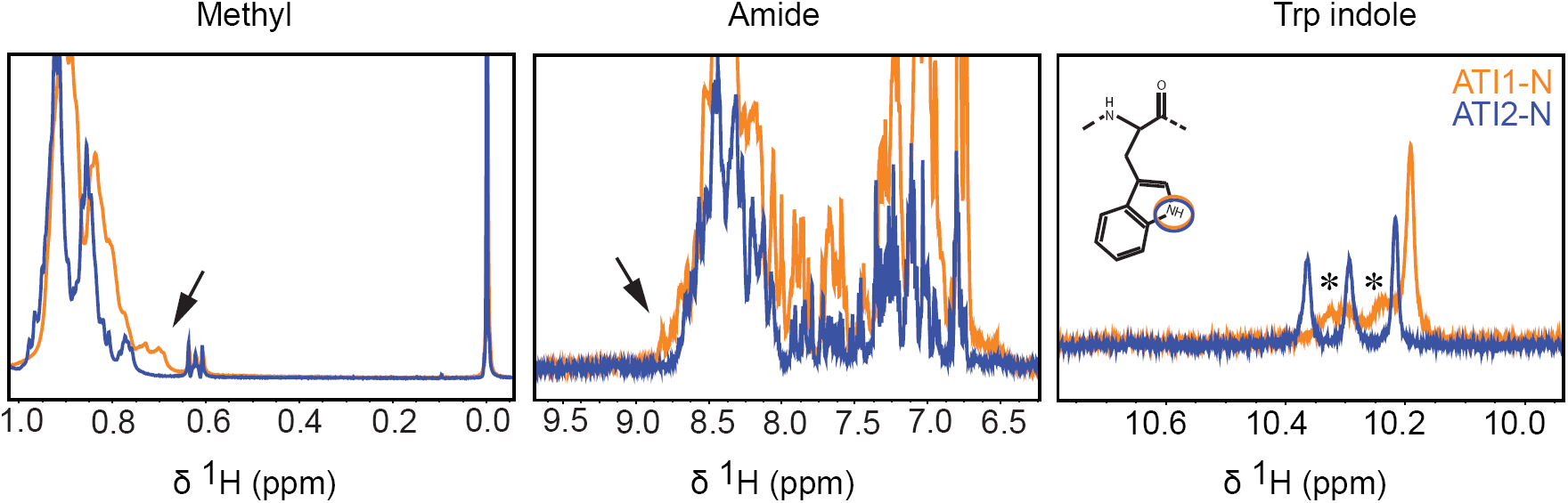
NMR spectra suggest a collapsed, pre-molten globule-like structure of ATI1-N, but a fully disordered ATI2-N. Zoom-ins on different parts of the 1D ^1^H NMR spectra recorded of ATI1-N and ATI2-N. Left, region of the spectrum with signals from methyl group protons (sharp peaks around 0.6 and 0.0 are from the 4,4-dimethyl-4-silapentane-1-sulfonic acid (DSS) reference); middle, region of the spectrum with signals from amide and aromatic protons; right, region of the spectrum with signals from Trp indole NH protons. Arrows indicate broader dispersion of signals of the methyl and amide protons in ATI1-N (orange) compared to ATI2-N (blue). Asterisks indicate broadening of the indole signals in AT11-N.

### The ATG8-interacting motif is present in the N-terminal IDRs of ATI1 and ATI2

ATI1 and ATI2 both contain potential AIMs in their N-terminal and C-terminal parts (Fig. 1A) [17]. The N-terminal AIMs have been proposed to be the functional sites of ATG8 interaction based on the fact that TM-HMM predicts the C-terminal part of ATI1 to be luminal [17], but no direct evidence supports this assertion. To test directly whether the N-terminal or C-terminal AIMs are functional, we first constructed point mutants in both putative AIMs in ATI2 and performed directed yeast two-hybrid assays with ATG8f. These experiments showed that ATI2 interaction with ATG8f was abolished upon mutation of the putative AIM in the N-terminal IDR, but not upon mutation of the putative AIM in the C-terminal part (Fig. 6A). Thus, the N-terminal, but not the C-terminal AIM of ATI2 is necessary for ATG8 interaction in yeast. We next used the well-resolved indole NH peaks in the 1D ^1^H NMR spectra to query whether the ATI1-N and ATI2-N were sufficient for ATG8 interaction: Spectra were recorded in the presence and absence of purified His_6_-ATG8f expressed in *E. coli* (Fig. 6B), and the regions of the spectra with chemical shifts from the indole NH protons were examined. This analysis showed clear line broadening of one specific indole NH signal in the presence of ATG8f, indicating that binding occurs to both ATI1-N and ATI2-N. We note that this positive binding result with ATG8f also validates the heterologously expressed ATI1-N and ATI2-N as functional proteins, despite their lack of a globular fold as shown in the preceding sections. We conclude that the N-terminal AIMs of ATI1 and ATI2 are necessary and sufficient for binding to ATG8f. We also conclude from the sharp peaks of the indole NH protons in the unbound state that the functional AIMs in both ATI1 and ATI2 reside in regions of high flexibility.

**Figure 6:**
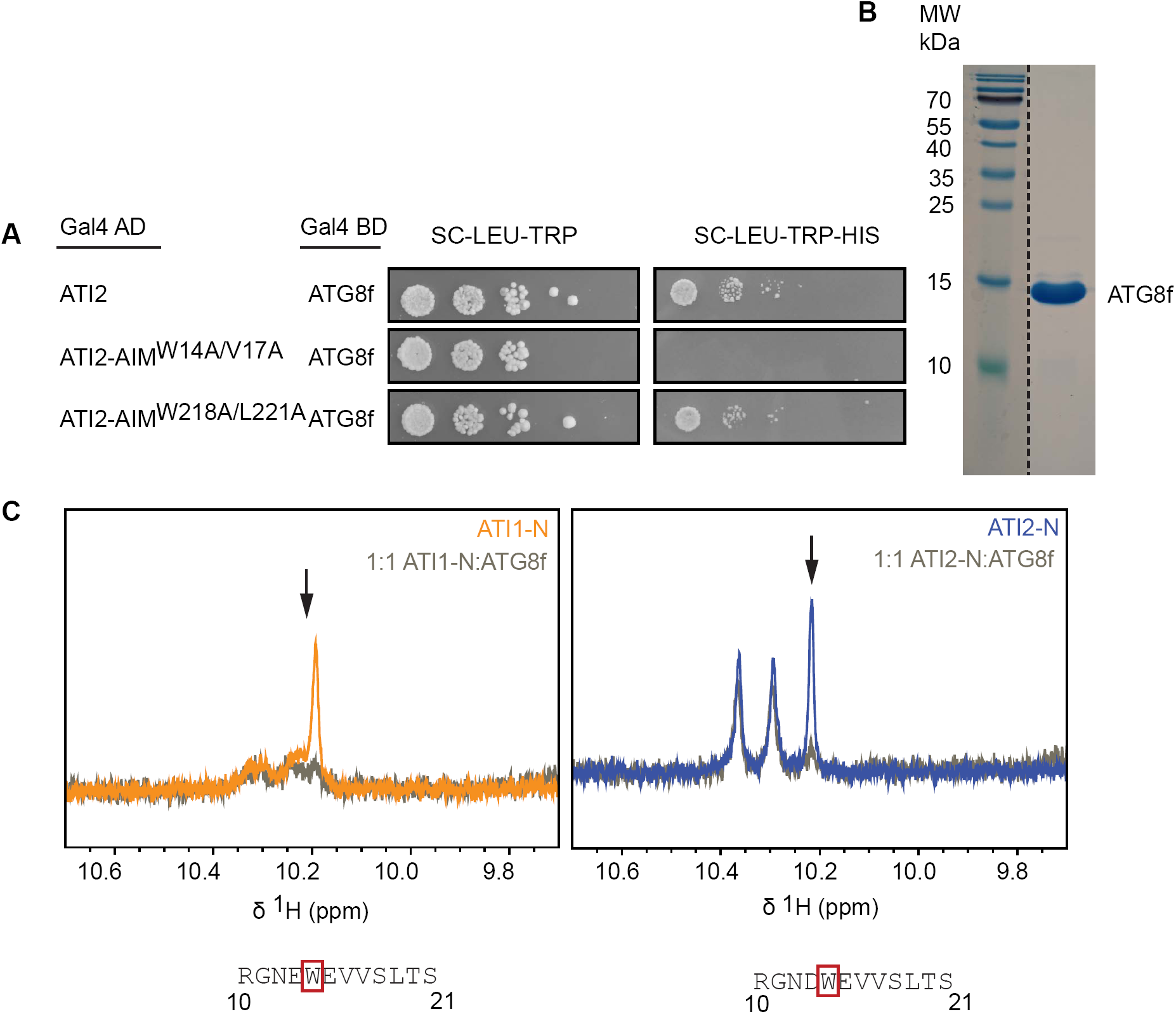
N-terminal AIMs are necessary and sufficient for ATG8 interaction. **A.** Yeast two-hybrid assay of interaction between ATI2 and ATG8f, including versions of ATI2 mutated in either the N-terminal putative AIM (ATI2-AIMW14A/V17A) or the C-terminal putative AIM (ATI2-AIM W218A/L221A). The *HIS3* auxotrophy marker is under Gal4 control, such that growth on –His plates is indicative of restoration of functional Gal4 by interaction between bait and prey fusions. *LEU2* and *TRP1* auxotrophy markers are expressed from the plasmids carrying the ATG8f-Gal4BD and ATI2-Gal4AD fusions, respectively. **B.** Recombinant His6-ATG8f expressed as the processed form (1-117) and subsequently purified by Ni^2+^affinity chromatography. Purity was checked on an 18 % denaturing polyacrylamide gel. **C.** Zoom-in on the region of the 1D ^1^H NMR spectra containing signals of the Trp indole NH. Spectra were recorded on ATI1-N (orange) and ATI2-N (blue) either alone or with a stoichiometric amount of purified ATG8f added to the sample (grey). The sequence of the N-terminal AIM is given below each spectrum, highlighting the Trp residue suggested to give rise to peak broadening upon ATG8f addition indicated by an arrow in the spectra.

### ATG8 interaction is coupled to post-translational modification of ATI1 and ATI2

One of the hallmarks of IDRs is their abundant post-translational modification that controls key aspects of their function *in vivo* [35]. The presence of post-translational modifications is often visible as mobility shifts compared to the unmodified protein in SDS-PAGE gels. We therefore raised ATI2 antibodies to see whether evidence of one or more forms could be discerned by simple western blotting. The antibodies were specific, because of the appearance of signals specific to wild type lysates as compared to lysates prepared from the mutant *ati2-2* carrying an exonic transposon insertion (Fig. 7A). Strikingly, ATI2 protein detected in microsomal fractions of wild type lysates migrated as two distinct species of roughly the same abundance, separated by a shift corresponding to 2-3 kDa. These different versions were not likely to be encoded by mRNAs arising from alternative transcription start sites, splicing events or polyadenylation sites: although inspection of RNA-seq [36] and cap analysis of gene expression (CAGE) data [37] revealed alternatively spliced transcripts (Fig. 7B), none of them could explain the presence of the observed equally intense ATI2 versions separated by 2-3 kDa (Fig. 7B, Fig. S1). By an analogous argument, only a single ATI1 protein is encoded by ATI1 mRNAs (Fig. 7B). Having established that at least one important function of the N-terminal IDRs of ATI1 and ATI2 is interaction with ATG8 through an AIM, we next constructed stable transgenic lines expressing 3xHA-tagged point mutants defective in ATG8 interaction. Comparison of the migration patterns of 3xHA-ATI1^WT^ and 3xHA-ATI2^WT^ with the corresponding AIM mutants showed a striking difference: In both cases, the AIM mutants ran at an apparent molecular weight slightly higher than the wild type proteins (Fig. 7C). This observation suggests that interaction of ATI1 and ATI2 with ATG8 is coupled to a change in their post-translational modification.

**Figure 7:**
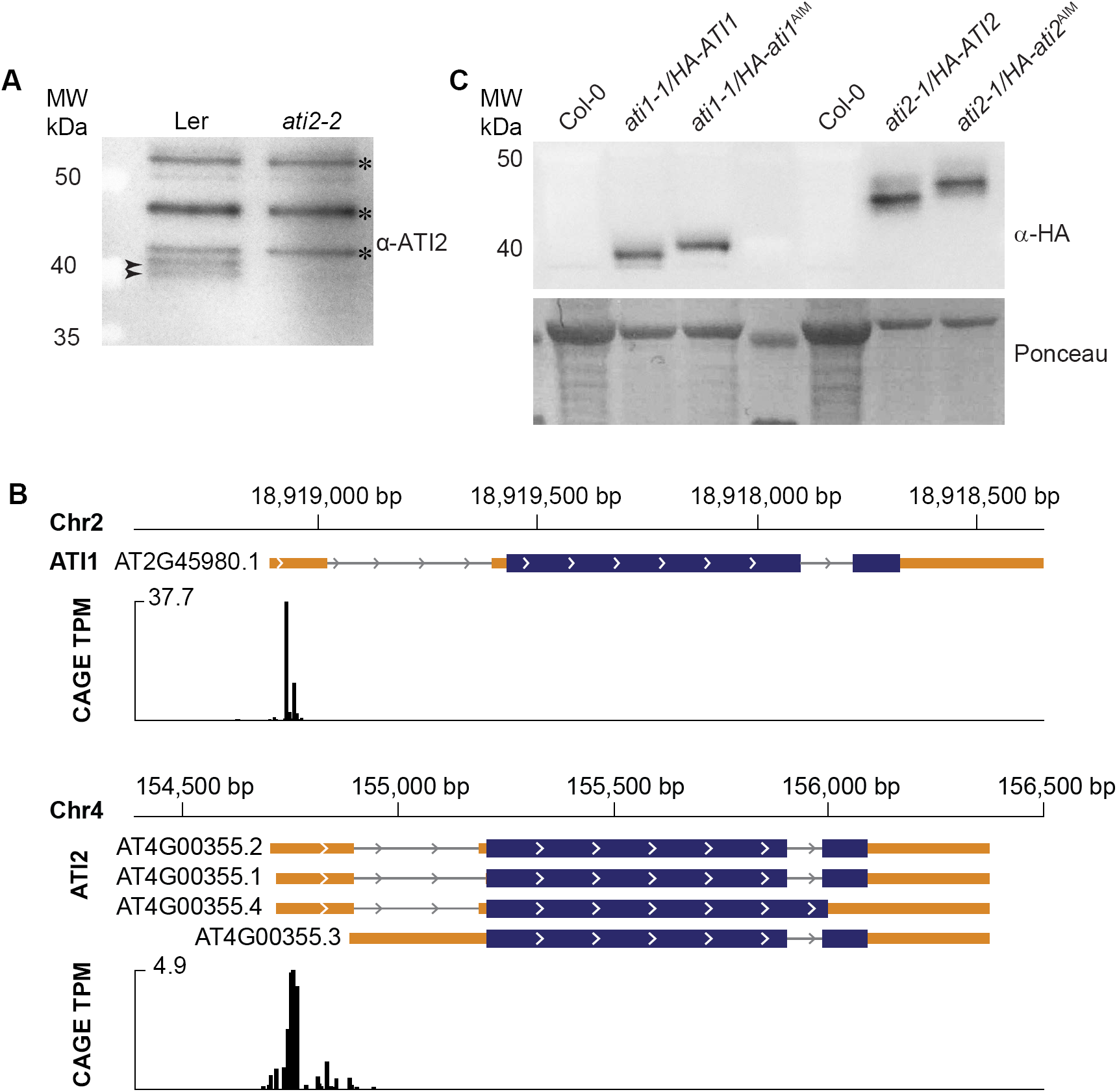
ATI1 and ATI2 are post-translationally modified depending on intact AIMs. **A.** Microsomal fractions isolated from Ler and *ati2-2* seedlings subjected to SDS-PAGE on a Bis-Tris 4-12% acrylamide gel and immunoblotted with anti ATI2 antibody. Arrows point to specific ATI2 bands whereas the asterisks indicate unspecific bands. **B.** Top, Araport11 [36] annotation of the *ATI1* gene and CAGE peaks mapping to *ATI1*. Bottom, same for the *ATI2* gene. CAGE data were taken from [37]. **C.** Total protein extracts of HA-ATI1, HA-ATI1AIM, HA-ATI2, and HA-ATI2AIM separated on a 10 % acrylamide gel and immunoblotted with anti HA antibody. Ponceau staining of the membrane shows loading and migration pattern of abundant proteins. Col-0 lysates were intentionally overloaded to show specificity of HA antibodies, and 2-fold less total protein of HA-ATI2 samples was loaded compared to HA-ATI1 samples to avoid obscuring the migration difference between HA-ATI2 and HA-ATI2^AIM^ by overexposure of the signals.

## DISCUSSION

### Direct experimental observation of structurally disordered AIMs in ATI1 and ATI2

Our biochemical and biophysical studies of ATI1 and ATI2 reveal several key properties of these two proteins crucial for a molecular understanding of their *in vivo* functions: We show that they are transmembrane proteins with long, intrinsically disordered N-terminal regions that contain the functional AIMs. Our study is, therefore, in line with the conclusion that functional AIMs occur in IDRs [7]. This conclusion was reached primarily based on prediction of disorder around many experimentally validated AIMs. Our study provides important experimental corroboration of this conclusion, because structural disorder of the Trp side chain engaged in ATG8 interaction was directly observed by NMR spectroscopy in the unbound state for both ATI1 and ATI2. Interestingly, clear structural differences between ATI1-N and ATI2-N were discernible from both size exclusion chromatography and NMR spectroscopy, with ATI1-N having a collapsed pre-molten globule-like structure with little secondary structure, while ATI2-N had properties of a fully disordered protein. Because of the similarity between the ATI1-N and ATI2-N sequences, it is of interest to identify exactly which protein properties cause this different behavior. It is of interest to consider whether this structural difference could have functional significance: in some instances, IDPs with residual helicity exhibit stronger binding to interaction partners than when their conformations are fully disordered [38,39]. If so, ATI1 and ATI2 may act in a sequential manner on *in vivo* cargoes such that ATI2 is mostly employed for degradation via selective autophagy at high cargo concentrations.

### Post-translational modification coupled to ATG8 interaction

Post-translational modification of cargo receptors or their cargoes is commonly employed to control the cargo-receptor interaction. For example, the interaction between DSK2 and BES1 is strongly enhanced upon DSK2 phosphorylation induced by stress [14], and GSNOR interaction directly with ATG8 is controlled via enhanced accessibility of its AIMs following hypoxia-induced *S*-nitrosylation of GSNOR [16]. Our observation that ATI1 and ATI2 mutated in the AIM migrate differently in SDS-PAGE than wild type ATI1 and ATI2 is distinct from these two examples, and has two important implications: First, it suggests that ATI1 and ATI2 are modified upon, rather than before, interaction with ATG8. Second, it may be a useful tool in future characterization of ATI1/2-cargo complexes, because it may allow easy distinction between ATG8-bound and free populations of these cargo receptors.

## MATERIALS AND METHODS

### Plant material

#### Plant material and growth conditions

Columbia-0 (Col-0) and Landsberg *erecta* (Ler) Arabidopsis thaliana accessions were used in this study. The *ati1-1* (SAIL404-D10, accession Col-0), *ati2-1* (GK142C08, accession Col-0) and *ati2-2* (GT_5_4264 JIC SM line, accession Ler) insertion lines were obtained from the Nottingham Arabidopsis Stock Centre. Seeds were sterilized as described [40] before being sown on Murashige-Skoog (MS) agar plates (4.3 g/L MS salts, 0.8% agar, 1% sucrose). Seedlings were grown for 14 days on MS at a constant temperature (21°C) and a 16-h light (80 µE m^-2^ s^-1^)/8-h darkness cycle.

#### Construction of transgenic lines

The coding sequences of *ATI1* and *ATI2* were amplified from genomic DNA with USER compatible primers (Table S1) and KAPA HiFi Hotstart Uracil + Readymix (Roche). The 3xHA tag was amplified from pSLF173 [41]. USER cloning was employed to combine and insert 3xHA/ATI1 PCR fragments into pCAMBIA2300u and 3xHA/ATI2 fragments pCAMBIA3300u [42]. The AIM of ATI1 was mutated (W14A and V17A) using USER compatible primers containing the mutations (Table S1) to amplify the fragments from the plasmid with *HA-ATI1*. *HA-ati1*^*AIM*^ was inserted into pCAMBIA2300U. The AIM of ATI2 was mutated (W14A/V17A) using site directed mutagenesis according to the Quickchange® protocol, Agilent Technologies, see Table S1 for primers used). Arabidopsis stable transgenic lines were generated by floral dip transformation [43] of *ati1-1* or *ati2-1* with *Agrobacterium tumefaciens* GV3101. Selection of primary transformants (T1) was done on MS-agar plates supplemented with kanamycin (50 µg/ml) for *HA-ATI1* and *HA-ati1*^*AIM*^ or glufosinate ammonium (PESTANAL; 7.5 µg/ml) for *HA-ATI2* and *HA-ati2*^*AIM*^. *HA-ATI1* and *HA-ati1*^*AIM*^ transgenic lines were analysed in the T2 generation, whereas *HA-ATI2* transgenic lines were T3 lines homozygous for transgene insertions. PCR was used to verify that the transgenic lines contained the correct transgenes (see Table S1 for genotyping primers), and all lines were analysed by western blots with HA antibodies to verify expression of the transgenes.

#### Microsome fractionation

Microsome fractionation was performed as described [40]. Briefly, 14-day old seedlings were flash frozen in liquid nitrogen and ground to a fine powder, and 1.2 mL of microsome buffer (50 mM MOPS, 0.5 M sorbitol, 5 mM EGTA, Roche protease inhibitors version 11 [one tablet/10 mL] at pH 7.6) was added to 0.2 g of ground tissue and vortexed thoroughly. Samples were spun at 8000*g* for 10 min at 4°C. Supernatants were transferred to new tubes and repeatedly spun at 8000*g* until no pellet was visible. Supernatants (“total extracts”) were spun at 100,000 *g* for 30 min at 4°C in a Beckman Optima XP ultracentrifuge. Pellets were resuspended in microsome buffer (50 mM MOPS, 0.5 M sorbitol, 5 mM EGTA, Roche protease inhibitors and repelleted by centrifugation at 100,000 *g* for 30 min at 4°C. Pellets were resuspended in 50 µl of microsome buffer, 1 M KCl, 0.1 M Na_2_CO_3_ or 1% sodium deoxycholate and incubated for 1h at 4 ?C. The samples were then spun at 100.000 *g* for 15 min at 4°C. Laemmli buffer was added to both the supernatant and pellet before boiling the samples and loading them on SDS-PAGE gels.

#### Antibodies and western blots

To raise antibodies against ATI2, two peptides corresponding to amino acid residues 35-46 (C+DSKDDDHKEVTP-CONH_2_) and 87-98 (C+KEGSDLSLKGLD-CONH_2_) were synthesized (Schafer-N, Copenhagen, Denmark), and a mix of the two peptides was used to immunize rabbits (Biogenes, Germany). For detection of 3xHA-ATI1 and 3xATI-ATI2, the monoclonal anti-HA antibody 12CA5 (1:2000, Roche) was used as the primary antibody, and anti-mouse HRP conjugated (1:5000, EMD Millipore) was used as the secondary antibody. For detection of PEPC, the polyclonal anti-PEPC antibody (1:2000, Rockland) was used as the primary antibody, and anti-rabbit HRP conjugated (1:10,000, Sigma) was used as the secondary antibody. To detect endogenous ATI2, the following procedure was used: Microsomal fractions were solubilized and boiled in Laemmli buffer and samples were loaded onto a 4-12% Bis-Tris Criterion™ acrylamide gel (Bio-Rad) and run in MOPS buffer (50 mM MOPS, 50 mM TRIS-Base, 0.1% SDS, 1 mM EDTA) pH 7.7. Proteins were blotted onto a nitrocellulose membrane and blocked for 5 min with 2 % BSA, washed three times in PBS-T buffer and incubated over night with anti ATI2 (1:500 in PBS-T with 2% BSA). The membrane washed then three times in PBS-T buffer and incubated with anti-rabbit HRP conjugated (1:10,000, Sigma) for 4 h. After an additional three washes in PBS-T, blots were developed using the enhanced chemiluminescent substrate [44] or SuperSignal West Femto Maximum Sensitivity Substrate (Thermo Fisher) and photographed with a SONY A7S camera.

#### RNA extraction, cDNA synthesis and Quantitative PCR (qPCR)

14-day old seedlings of Col-0 were flash frozen in liquid nitrogen and grounded to a fine powder. 100 mg was vortexed in 1 ml of TRI Reagent (Sigma) and 200 µl chloroform was added. Total RNA was precipitated from the aqueous phase with isopropanol and the pellet was washed twice with 70% EtOH. The pellet was resuspended in water. 1 µg of RNA was reverse transcribed into cDNA using oligo dT primers and the SuperScript™ III kit according to the manual. qPCR reactions were made with Maxima SYBR Green qPCR Master Mix (Thermo Scientific) and run on a BioRad CFX connect thermal cycler using primers designed to distinguish the four isoforms of Col-0 *ATI2* and *ACT2* as the control (Table S1). The expression levels of the *ATI2* isoforms were presented as cycle threshold (Ct) relative to *ACT2* as an average of three technical replicates with standard deviation as error bars.

#### Expression and purification of recombinant ATI1-N and ATI2-N

The N-terminus of *Arabidopsis thaliana* ATI1 (aa 1-180, At2G45980) and ATI2 (aa 1-193, At4G00355) were amplified from full-length coding sequences of ATI1 and ATI2 cloned into pGEMTeasy, using oligonucleotides 1/2 and 3/4 (Table S1). PCR fragments were ligated into pCR™4/TOPO™ using the TOPO™ TA Cloning™ Kit for Sequencing (Invitrogen). ATI1-N and ATI2-N were excised and ligated into the expression vector pGEX4-T1 where they were N-terminally fused to a glutathione-S-transferase (GST)-tag downstream of an introduced tobacco etch virus (TEV) cleavage site (ENLYFQS). Recombinant protein was expressed in *E. coli* BL21 (DE3 7tRNA) Rosetta cells grown in 1 liter Luria Bertani (LB) media supplemented with ampicillin (100 µg/ml) and chloramphenicol (33 µg/ml) to an OD_600_ = 0.6-0.8 before protein expression was induced by 1 mM isopropyl β-D-1-thiogalactopyranoside (IPTG) at 18 °C O/N. Cells were harvested and lysed by 2-3 passages through a French press (≥10,000 psi) in freshly prepared cold lysis buffer (1x PBS, 1mM tris (2-carboxyethyl) phosphine hydrochloride (TCEP), 1 complete EDTA-free protease inhibitor tablet (Roche)). Cell lysate was centrifuged at 12,500 *g* for 1 hour at 4 °C and the supernatant was filtered through a 0.45 µm filter. The cleared supernatant was incubated on glutathione-conjugated sepharose beads (GE Healthcare) for 1 hour at 4 °C. The beads were washed with cold 1x PBS and the GST-tag was cleaved off O/N at ∼18 °C by His_6_-TEV protease (estimated 1:30 molar ratio) made in house in 50 mM tris-HCl pH 7.2, 50 mM NaCl, 0.5 mM EDTA, 1.75 mM TCEP. The flow through was collected, spun down and His_6_-TEV was captured on Ni-NTA beads (Macherey-Nagel) for 1 hour at 4 °C. ATI1-N was concentrated on PES membrane Vivaspin® 500 columns (Sartorius). Precipitated protein was dissolved in 6 M urea, 2% sodium dodecyl sulphate (SDS), 10 % glycerol, 50 mM Tris-HCl pH 6.8, pooled with the soluble fraction and dialyzed (ZelluTrans dialysis tubes, 6,000-8,000 MWCO (Roth)) O/N into the buffer used for size exclusion chromatography (20 mM tris-HCl pH=7.2, 5 mM NaCl, 1 mM β-mercaptoethanol). ATI2-N was buffer-exchanged into the size exclusion chromatography buffer by means of PES membrane Vivaspin® 500 columns (Sartorius).

#### Expression and purification of recombinant ATG8f

The processed form of *Arabidopsis thaliana* ATG8f (aa 1-117, At4G16520.1) was amplified from a full length pCR8-ATG8f cDNA clone using oligonucleotides 5/6 (Table S1) and ligated into the pCR™4 TOPO™ vector (Invitrogen). ATG8 was excised and ligated into the expression vector pET15b where it was N-terminally fused to a His_6_-tag downstream of an introduced TEV protease cleavage site (ENLYFQS). Recombinant protein was expressed in *E. coli* BL21 (DE3 7tRNA) Rosetta cells grown in 2 litre LB media supplemented with kanamycin (50 µg/ml) and chloramphenicol (33 µg/ml) to an OD_600_ of 0.6-0.8 before protein expression was induced by 0.5 mM IPTG at 18 °C O/N. Cells were harvested and lysed by 3 passages through a French press (≥10,000 psi) in cold lysis buffer (20 mM tris-HCl pH=8, 10 mM imidazole, 300 mM NaCl, 1 mM TCEP, 1 complete EDTA-free protease inhibitor tablet (Roche)). Cell lysate was centrifuged at 30000 *g* for 30 min at 4 °C and supernatant was filtered through a 0.45 µm filter. The cleared supernatant was incubated on immobilized nickel-chelating nitrilotriacetic acid (NTA) agarose beads (Macherey-Nagel) for 2 hours at 4 °C. The beads were washed with cold 20 mM imidazole, 20 mM Tris-HCl pH 8, 300 mM NaCl and the His_6_-tag was cleaved off O/N at 18°C by His_6_-TEV protease (estimated ∼1:80 molar ratio) made in house in 50 mMTtris-HCl pH 7.2, 50 mM NaCl, 0.5 mM EDTA, 1.75 mM TCEP. The ATG8f-containing flow through was collected and dialyzed (ZelluTrans dialysis tubes, 6,000-8,000 MWCO (Roth)) into 20 mM 2-[N-morpholino] ethanesulphonic acid (MES)-KOH pH=6 O/N at 4 °C. The dialyzed ATG8f-containing sample was loaded onto a methyl sulfonate MonoS 5/50 GL cation exchange column (GE Healthcare) connected to an HPLC Äkta Purifier system (GE Healthcare). Protein was eluted by gradually increasing the ionic strength of the mobile phase to 1 M NaCl in 20 mM MES-KOH pH 6. Eluate was monitored at A_280_ and purity checked by SDS PAGE analysis. ATG8f-containing fractions were buffer exchanged into ddH2O using 10,000 MWCO PES membrane Vivaspin® 500 columns (Sartorius) and subsequently flash frozen in liquid nitrogen and lyophilized.

#### Yeast two-hybrid analysis

The yeast two hybrid analysis was performed using the Clontech Gal4 Matchmaker system with bait vector pGBKT7 and prey vector pGADT7. ATG8f cDNA was amplified from pCR8-ATG8f (see above) and cloned into pGBKT7, while ATI2 and was cloned into pGADT7 (see Table S1 for primers used). The ATI2-AIM mutants were generated by site-directed mutagenesis of the pGADT7-ATI2 plasmid using the Quickchange protocol and primers listed in Table S1. Bait and prey plasmids were co-transformed into the *S. cerevisiae* reporter strain PJ694A according to Invitrogen instructions (Matchmaker™ GAL4 Two-Hybrid System 3 & Libraries User Manual, Protocol No. PT3247-1 Version No.PR742219) and using Invitrogen dropout media. Interaction tests by spotting assays were done in the following way: Freshly transformed colonies were restreaked on –Trp,Leu plates and grown at 30°C for two days. Colonies were then resuspended in water, and OD_600_ was adjusted to 3.0 for all strains. Five µl of 10-fold serial dilutions were then spotted on either dropout medium without Trp and Leu to control for strain growth, or dropout medium without Trp, Leu and His and supplemented with 1.5 mM 3-amino-1,2,4-triazole (Sigma) to test interaction between bait and prey fusions to individual Gal4 domains. Pictures were taken after incubation for two days at 30°C.

#### Size exclusion chromatography of ATI1-N, ATI2-N and prolactin

ATI1-N, ATI2-N and human prolactin (hPRL) purified as described [45] were subsequently loaded onto a HiLoad Superdex™ 16/600 200 prep grade column (GE Healthcare) connected to an HPLC Äkta Purifier (GE Healthcare) equilibrated with 1.5 column volumes of the size exclusion chromatography buffer (20 mM Tris-HCl pH 7.2, 5 mM NaCl, 1 mM β-mercaptoethanol). Eluting protein was monitored at A_280_.

#### Heat aggregation test of ATI1-N and ATI2-N

In two parallel setups, equal volumes of ATI1-N, ATI2-N and GST were heated to 75°C for 10 min or to 95°C for 5 min. Samples were spun down and aliquots were taken from the supernatant and the pellet for subsequent SDS-PAGE analysis. 4x lithium dodecyl sulphate (LDS) sample buffer was added to samples before they were boiled at 75 °C for 5 min and subsequently spun down. Samples were loaded on a 14 % SDS PAGE and run in 1x Tris-glycine buffer, 0.1% SDS. Separated protein was stained in Instant Blue (Expedeon).

#### Circular dichroism spectroscopy of ATI1-N and ATI2-N

Circular dichroism measurements of ATI1-N and ATI2-N were obtained on a Jasco J-810 spectropolarimeter equipped with a Peltier temperature controller. Protein was measured in a 1 mm path length quartz cuvette at 20°C. 5 uM protein was measured in 10 mM NaH_2_PO_4_ pH 7.2, 20 mM Na_2_SO_4_. All runs were obtained with a scan speed of 50 nm/min, a step size of 0.5 nm and a bandwidth of 1 nm. Spectra were acquired as 10 accumulations from 260-198 nm, subtracted buffer absorption recorded at identical settings and Fourier transformed utilizing the software Spectrum Analysis. Data was subsequently converted from mdeg to molar ellipticity normalized to molar concentration of peptide bonds according to

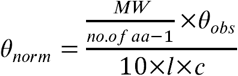

Where MW is the molecular weight in Da, θ_obs_ is the observed ellipticity in mdeg, l is the pathlength in cm, and c is the concentration in g/l. The resulting unit is mdeg cm^2^ dmol^-1^.

#### NMR spectroscopy of unlabeled ATI1-N and ATI2-N without and with the addition of ATG8f

Expressed and purified ATI1-N (270 uM) and ATI2-N (140 uM) were prepared individually in buffer resembling physiological conditions (20 mM Tris-HCl pH 7.2, 150 mM KCl, 5 mM NaCl, 1 mM MgCl_2_, 3 mM TCEP) containing 10 % D_2_O and 250 uM 4,4-dimethyl-4-silapentane-1-sulfonic acid (DSS) for referencing. Samples with stoichiometric amounts of ATI1-N and ATG8f (70 uM) and ATI2-N and ATG8f (70 uM) were prepared in the same buffer containing 250 uM DSS and 10 % D_2_O. 100-120 ul the samples were applied to a 3 mm Shigemi NMR tube. The 1D ^1^H spectra were recorded on a Bruker Avance III 600 MHz spectrometer equipped with a TCI cryoprobe, using pulse sequences as described [46]. Bruker TopSpin 3.5 (pl5) software was used to control the spectrometer and to process the spectra.

## ACKNOWLEDGEMENTS

This work was supported by the Novo Nordisk Foundation (Hallas Møller Stipend 2010, to PB), the European Research Council (ERC-StG Micromecca 282460, to PB), and the Danish Research Councils (to BBK, 4181-00344). Villumfonden is thanked for a generous NMR instrument grant. Axel Thieffry is thanked for analysis of transcription start site data for *ATI1* and *ATI2*. Theo Bølsterli and his team are thanked for plant care.

